# Reaction Pathway Sampling and Free Energy Analyses for Multimeric Protein Complex Disassembly with Employing Hybrid Configuration Bias Monte Carlo/Molecular Dynamics Simulation

**DOI:** 10.1101/2020.09.16.299263

**Authors:** Ikuo Kurisaki, Shigenori Tanaka

## Abstract

Physicochemical characterization of multimeric biomacromolecule assembly and disassembly processes is a milestone to understand the mechanisms for biological phenomena at molecular level. Mass spectroscopy (MS) and structural bioinformatics (SB) approaches have become feasible to identify subcomplexes involved in assembly and disassembly, while they cannot provide atomic information sufficient for free energy calculation to characterize transition mechanism between two different sets of subcomplexes. To combine observations derived from MS and SB approaches with conventional free energy calculation protocols, we here designed a new reaction pathway sampling method with employing hybrid configuration bias Monte Carlo/Molecular Dynamics (hcbMC/MD) scheme and applied it to simulate disassembly process of serum amyloid P component (SAP) pentamer. The results we obtained are consistent with those of the earlier MS and SB studies with respect to SAP subcomplex species and the initial stage of SAP disassembly processes. Furthermore, we observed a novel dissociation event, ring-opening reaction of SAP pentamer. Employing free energy calculation combined with the hcbMC/MD reaction pathway trajectories, we moreover obtained experimentally testable observations on (1) reaction time of the ring-opening reaction and (2) importance of Asp42 and Lys117 for stable formation of SAP oligomer.

## Introduction

The physicochemical entity of various biological phenomena is multimeric biomolecule complexes formed transiently or stably in the cell. The molecular components of such complexes have been extensively identified and individually assigned to corresponding biological functions (*e.g*., signal transduction^1–3^, DNA repairs^4, 5^, protein systhesis^6, 7^, and RNA splicing^8, 9^). Since these functions are expressed and regulated through biomolecule assembly and disassembly in response to the cellular environment^10–12^, physicochemical characterization of these processes is a milestone to understand the mechanisms for biological phenomena at molecular level.

Multiple biomacromolecule assembly and disassembly processes can be physicochemically characterized by solving the two essential problems coupled with each other: (1) identifying a set of intermediate states characterized by subcomplexes appearing in these processes; (2) elucidating physicochemical mechanisms of transition between such intermediate states.

Mass spectrometry (MS) analyses have become a powerful tool to systematically analyze relative subunit-interface strength and provide a possible set of subcomplexes undergoing assembly and disassembly processes at molecular level.^13–15^ Meanwhile assembly and disassembly orders have been extensively studied in the field of structural bioinformatics (SB) during the last fifteen years^16^ and can now be predicted with high accuracy based on atomic structures of individual subunits.^17^ Thus identification of subcomplexes involved in assembly and disassembly processes has become technically feasible, then providing important clues to characterize intermediate states consisting of assembly and disassembly processes.

As for physicochemical analyses of mechanisms for transition between such intermediate states, atomistic simulations are practically useful at present; we could promptly find a feasible approach for the problem in a class of conventional free energy profile (FEP) calculation protocols, *e.g*., umbrella sampling combined with reaction pathway sampling techniques^18, 19^ FEP such as potential of mean force (PMF) can provide quantitative insights into physicochemical mechanisms for biomacromolecule complex formation in terms of evaluated rate constants of configurational transition between two different intermediate states. Then it could seem practically feasible to examine physicochemical mechanisms for multimeric biomacromolecule assembly and disassembly processes.

Nevertheless, there still remains one technical problem upon combining the subcomplex identification methods with FEP calculation protocols. They need *a priori* knowledge of a complete set of atomic coordinates for all subunits and subcomplexes at intermediate states in assembly or disassembly processes.

In fact, it is still challenging for the above mentioned state-of-the-art MS and SB approaches to provide atomic information sufficient for FEP calculation protocols. MS approaches identify subunits and subcomplex species as particles with molecular mass and net charge, so that their spatial resolution is bound to macromolecular level. SB approaches directly predict a final docking pose and a pair of dissociated subcomplexes. It is therefore out of their methodological design to identify a configuration of a complete set of subunits, at each intermediate state involved in a pre-binding and post-unbound conditions.

To solve the technical problem in molecular simulation, we here propose a new reaction pathway sampling method. The theoretical framework we here employ is a hybrid Monte Carlo (MC)/Molecular Dynamics (MD) scheme, since configuration sampling method in the MD phase is arbitrarily selected for specific research purposes, then resulting in flexible applications of this simulation framework to a variety of molecular phenomena in condensed phases^20–23^.

We can find research purpose-oriented flexibility within magnificent examples of hMC/MD simulation studies. Chemical bond formation and scission are empirically simulated by switching potential functions and using effective potential function in the frameworks of REDMOON^21^ and KIMMDY^23^, respectively. Meanwhile TRS^20^ and hybrid neMD-MC^22^ employ randomly assigned force bias and perturbed potential energy to accelerate transformation of biopolymer conformation, respectively.

In the present method, we employed steered molecular dynamics (SMD) method to enhance non-equilibrium subprocesses composing assembly and disassembly processes, namely, inter-subunit association and dissociation reactions. This method have been widely used to simulate these reactions involved in dimeric biomacromolecule complexes.^19, 24, 25^ Aiming additional enhancement of configuration sampling, we applied the configuration bias Monte Carlo (cbMC) scheme^26–28^ to hybrid MC/MD simulation procedure, recalling the earlier hybrid MC/MD study resulting in several time enhancement of lipid bilayer configuration sampling^29^.

To test our simulation method, referred to as hybrid configuration bias MC/MD (hcbMC/MD) method hereafter, we initially considered disassembly processes because of technical straightforwardness. The hcbMC/MD method is applicable to simulate disassembly processes with simply employing inter-subunit dissociation reactions without subunit-subunit docking pose prediction which is possibly needed to examine assembly processes associated with multiple subunits. In this study, the hcbMC/MD method was applied to simulate disassembly processes of Serum Amyloid P component (SAP) homo pentamer (Figure 1), whose subcomplex species appearing in disassembly processes have been extensively studied in the previous MS studies.^13, 30, 31^.

**Figure 1.**
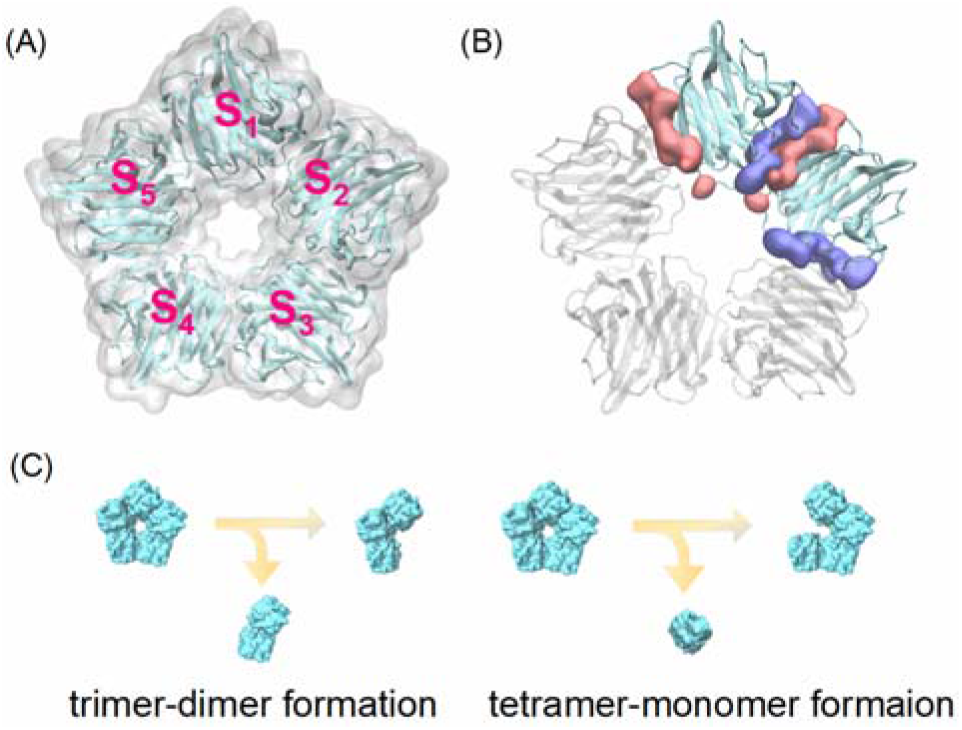
Molecular structures of serum amyloid P component (SAP) homo pentamer. (A) SAP pentamer with annotation for each subunit. (B) Two individual inter-subunit interaction surfaces of SAP subunit, highlighted by blue and red colors; S_1_ and S_2_ are depicted as representative of subunit pair. (C) Initial stages of SAP pentamer disassembly processes predicted by structural bioinformatics approach^13^. Reactions shown in panel C are supposed for any possible monomer and dimer.

## Materials and Methods

### Setup of serum amyloid P component system

We used atomic coordinates of serum amyloid P component (SAP) homo-pentamer (PDB entry: 4AVS)^32^. Two calcium ions bound to SAP subunits were retained in the atomic coordinates of this SAP system. Nε protonation state was employed for each of histidine residues, and all carboxyl groups in aspartate and glutamate residues were set to the deprotonated state, where we considered physiological pH condition in the cell. The SAP structure was solvated in the rectangular box with 49305 water molecules and was electrically neutralized by adding 10 Cl^-^ ion molecules.

To calculate the forces acting among atoms, AMBER force field 14SB^33^, TIP3P water model^34, 35^, and JC ion parameters adjusted for the TIP3P water model^36, 37^ were employed for amino acid residues, water molecules, and ions, respectively. In addition, the force field parameter sets developed by Bradbrook was applied for divalent calcium ion^38^. Molecular modeling of the SAP system was performed using the LEaP modules in AmberTools 17 package^39^. Molecular mechanics (MM) and molecular dynamics (MD) simulations were performed under the periodic boundary condition with GPU-version PMEMD module in AMBER 17 package^39^ based on SPFP algorithm^40^ with NVIDIA GeForce GTX1080 Ti. The solvated structure was relaxed through 10-ns NPT MD simulations, then being employed as the initial atomic coordinates for the following hybrid configuration bias MC/MD simulations. Further computational details are described in **S-1** and **S-2** in Supporting Information.

### Design concept of hcbMC/MD-based path sampling

Our path sampling simulation method is designed while considering the three essential factors which possibly have prevented MD simulations from their application to the assembly and disassembly processes associated with multiple biomacromolecules: (1) physicochemical events rarely occurring within available computational time; (2) technical complexity to prespecify a sequence of intermediate states, whose possible candidates increase with factorial of the number of inter-subunit interface; (3) no *a priori* knowledge for a full set of atomic coordinates of macrobiomolecules at each intermediate state in assembly and disassembly processes.

We cope with the first factor by using two theoretical techniques: (1) steered MD (SMD) method to enhance diffusion of subunit molecules in each MD phase; (2) a configuration bias Monte Carlo scheme in each MC phase (*vide infra*).

Assembly and disassembly processes are sequences of inter-subunit association and dissociation reactions, respectively. Even such an elementary subprocess generally takes several micro seconds or longer^41–43^, thus rarely occurring within brute-force MD simulations available of today. Enhanced sampling such as SMD method is indispensable to simulate such rare events within realistic computation time^19^.

Besides aiming to further accelerate development of molecular systems toward disassembly, we employed biased configuration selection^26–29^ for a set of generated configurations by using a purpose-oriented biased potential energy function, as shown below.

Meanwhile a configuration generated by enhanced sampling is rejected with certain probability by Metropolis algorithm in MC phase. Acceptance judgement of generated configurations via Metropolis algorithm is implemented to circumvent selection of subcomplex configurations anomalously deformed by applying external forces.

The second factor is treated by using a set of the predefined reaction coordinates, where we recall successful usage of predefined set of chemical reactions in REDMOON method^44^. At every hcbMC/MD simulation cycle, a reaction coordinate for inter-subunit association and dissociation is randomly chosen from a set of the predefined reaction coordinates and the system is propagated in MD phases (*see* Materials and Methods section for predefined reaction coordinates specific for the SAP system below). Random choice of reaction coordinates is implemented to circumvent technical difficulty to prespecify a complete sequence of inter-subunit association and dissociation reactions. As for the case of SAP system, we can simulate the disassembly process without selecting probable sequences of dissociation reactions among the 5! (= 120, derived five inter-subunit interface of SAP pentamer) candidates.

Employing a predefined set of reaction coordinates in enhanced sampling simulations coincidently provides a solution for the third factor; it enables us automatically search reaction pathways without *a priori* knowledge of a complete set of atomic coordinates of all subunits and subcomplexes in molecular system at each intermediate state in the assembly and disassembly processes. In application to disassembly processes, our pathway sampling method needs only initial atomic coordinates of molecular system, which can be obtained from structural biological techniques (*e.g*. X-ray crystallography, electron microscopy and so on). This point differs from the conventional ones, such as Nudged Elastic Bond^45^ and targeted MD^46^ methods, which at least require a set of atomic coordinates of molecular configurations with respect to both the initial and final states.

It is worthwhile to note that multi-trajectory approaches to aim rare event sampling, *e.g*., Weighted Ensemble (WE) method^47^, could be potentially applicable to solve the problems we are addressing in this study. Nonetheless, considering available computational resources of today, it still seems to be challenging for atomistic WE simulations to examine assembly/disassembly processes of multimeric biomacromolecule complexes at atomic level. Detailed discussion is given in Supporting Information.

### Hybrid configuration bias MC/MD simulation scheme

The simulation scheme is designed based on a general configuration bias MC method^26^ as follows. We here employed unbiased and SMD simulations to generate old and new sets of atomic configurations of molecular systems, respectively. Inter-subunit dissociation reactions are mainly accelerated with SMD simulations. Detailed simulation conditions of MD and SMD-based configuration generation are given in the next subsection.

1. Generate *M* trial configurations {**a**_1_, **a**_2_,⋯**a**_*M*_} by an unbiased MD simulation and calculate bias energy *u^bias^*, which will be specified later, for each configuration.
2. Assign the last snapshot obtained from the simulation (**a**_*M*_) as an old configuration, denoted by **x**_*o*_ and define the Rosenbluth factor^26^

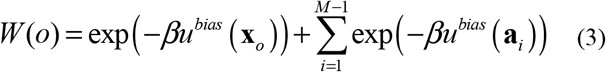

*β* denotes inverse of k_B_T, where k_B_ and T are Boltzmann constant and system temperature, respectively. In the following step 3, new configurations are generated by starting from **a**_*M*_, so that we assigned this configuration to **x**_*o*_ by recalling conventional cbMC schemes^26^.
3. Generate *M* trial configurations {**b**_1_, **b**_2_,⋯**b**_*M*_} by a steered MD simulation starting from the old configuration **x**_*o*_ and calculate bias energy *u^bias^* for each configuration.
4. Define the Rosenbluth factor

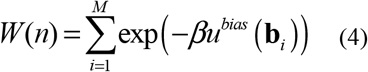

and select one configuration among {**b**_1_, **b**_2_, ⋯ **b**_*M*_}, denoted by **x**_*n*_, with a probability

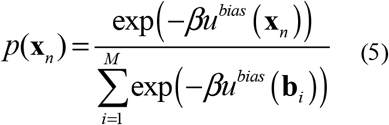
5. The configuration change is accepted with a probability

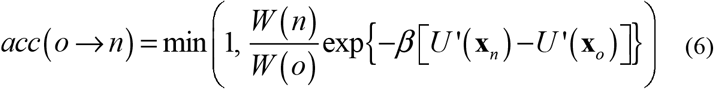

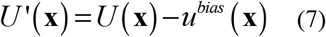

where **x** and *U*(**x**) denote a configuration and the potential energy function of the system with configuration of **x**, respectively.

Considering that configurations of the system are generated by moving all atoms simultaneously with molecular dynamics method, we used the Rosenbluth factor in the forms given in Eqs. (3) and (4). Monte Carlo trial with this type of configuration sampling is particularly referred to as orientation bias Monte Carlo^26^.

As for the right-hand side of Eq (4), we assume that probability of generating forward move (**x**_*o*_ → **x**_*n*_) is equal to that of generating backward move (**x**_*o*_ ← **x**_*n*_). This assumption is employed here for simplification of simulation implementation but not trivial in general. Our hcbMC/MD method could work for extensively sampling configuration space, while it is uncertain to exactly construct a canonical ensemble defined by an unperturbed Hamiltonian. Then with the aim of validating physicochemical observations obtained from an hcbMC/MD trajectory, it is a practical option to perform additional umbrella sampling MD simulations with a set of snapshot structures involved in an inter-subunit dissociation event during the trajectory.

A multimeric protein complex disassembly process should be accompanied by gradual breakage of inter-subunit native contacts (NC). Then we defined the bias energy as a function of the total number of inter-subunit NC formed in a multimeric protein complex (*n_NC_*), where the initial atomic coordinates are used as the reference structure:

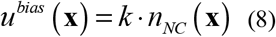

The coefficient *k* is set to *β*^−1^ at 300 K, that is 0.6 [kcal/mol] by assuming energetic stabilization obtained from hydrophobic atomic contacts^48^. A set of native contacts at subunit binding interface was defined by using the initial atomic coordinates of SAP pentamer for hcbMC/MD simulations. All hcbMC/MD simulations were performed with employing an in-house MC/MD simulation interface.

### Configuration generation at each hcbMC/MD cycle by using unbiased and steered MD simulations

Configurations of SAP system, denoted by {**a**_1_,**a**_2_,⋯**a**_*M*_}, were sampled by using an unbiased NPT MD simulation (300 K, 1 bar). Meanwhile, configurations of the system undergoing a partial inter-subunit dissociation, denoted by {**b**_1_, **b**_2_, ⋯**b**_*M*_}, were sampled by using a steered molecular dynamics (SMD) simulation under the NPT condition (300 K, 1 bar), starting from the snapshot structure obtained from the unbiased MD simulation. The simulation length is 100 ps for each of MD and SMD simulations. A set of atomic coordinates was recorded by every 10 ps and was indexed in ascending order to prepare {**a**_1_, **a**_2_,⋯**a**_*M*_} and ·**b**_1_, **b**_2_,⋯**b**_*M*_}; M is set to 10 in this simulation condition. The system temperature and pressure were regulated by Langevin thermostat with 1-ps^-1^ collision coefficient, and Monte Carlo barostat with attempt of system volume change by every 100 steps, respectively.

In every hcbMC/MD cycle, we randomly chose a pair of monomers in the 10 (= _5_C_2_, derived from combination of two subunits among the five) candidates. We employed the distance between centers of gravity of the chosen subunit pair as the reaction coordinate of SMD simulation. A center of gravity for a SAP subunit was calculated by using all of the 204 C_α_ atoms.

In an SMD simulation, a target value of the distance was set to *d*_0_ + Δ*d*, where *d*_0_ and Δ*d* are an initial value of the distance at the cycle and a random integer in the range of 8 to 12, respectively. The harmonic potential with the force constant of 10 kcal/mol/Å^2^ was imposed on the reaction coordinate. In each cycle, a reaction coordinate for inter-SAP subunit dissociation reaction and a random seed for Langevin thermostat were changed randomly. Following the above simulation conditions, an unbiased MD simulation was also designed to evaluate the sampling performance of hcbMC/MD simulation, where the 100-ps MD simulation cycle was repeated 2000 times. Additional computational details and a corresponding unbiased MD simulation procedure are described in **S-1** and **S-3** in Supporting Information, respectively.

### Umbrella sampling molecular dynamics simulations

Certain numbers of snapshot structures were extracted from a hcbMC/MD trajectory and employed for USMD windows. Following temperature relaxation simulations, 5-ns NVT USMD simulations (300 K) were performed for each of the USMD windows (*see* Table S1 in Supporting Information for details). The system temperature was regulated using Langevin thermostat with 1-ps^-1^ collision coefficient. Each of the last 3-ns USMD trajectories was used to construct a PMF (*see* Figure S1 for convergence of PMF curves). A reaction coordinate is the distance between centers of gravity of the subunit pair undergoing dissociation reaction. A set of USMD simulations was repeated by 8 times. Besides, we calculated PMFs of stable SAP pentamer with employing the initial atomic coordinates for the hcbMC/MD simulations. The further technical details are given in Supporting Information (*see* **SI 4** and **SI 5**).

### Trajectory analyses

Root mean square deviation (RMSd), and native and non-native atomic contacts were analyzed with the cpptraj module in AmberTools 17 package^39^. We calculated RMSd and native contacts by using the C_α_ atoms and the non-hydrogen atoms in the initial atomic coordinates for the hcbMC/MD simulations, respectively. The distance criterion for atomic contacts was set to 3.5 Å. Inter-subunit dissociation and reassociation conditions are defined as follows: N_native_ = 0 and N_non-native_ = 0; N_non-native_ ≥ 0 in N_native_ > 0, respectively. N_native_ and N_non-native_ denote the number of native atomic contacts and that of non-native atomic contacts made between a pair of SAP subunit, respectively. Collision cross section (CCS) of SAP protein was analyzed by using IMPACT program^49^.

PMF was calculated with Weighed Histogram Analysis Method (WHAM)^50, 51^ by using each set of USMD trajectories. Statistical errors of PMF values, ***σ***_*PME*_(*ξ*), were estimated by employing bootstrapped sampling^52^:

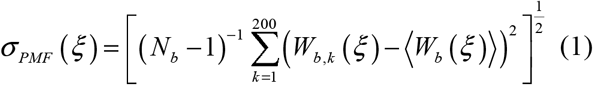

Here, *N_b_, ξ*, and *W_b,k_*(*ξ*) denote the number of bootstrapped sampling, the reaction coordinate and the value of k^th^ bootstrapped PMF at each point of *ξ*, respectively. 〈*W_b_*(*ξ*)〉 is the average over all *W_b,k_* (*ξ*), where *k* ranges from 1 to 200.

Reaction rate, *k_TST_*, is estimated by using Eyring’s transition state theory:

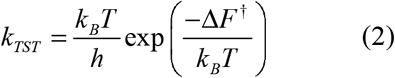

where *k_B_* and *T* denote an activation barrier height, Planck constant, Boltzmann constant and a temperature of system, respectively. The estimation with employing formula (2) is supposed to be an upper bound of the reaction rate.^53^

Reaction time scale, *τ*_TST_, is defined as the inverse of *k_TST_*. Δ*F*^†^ is defined as *F*(*ξ*_0_′)–*F*(*ξ*_0_), where the value of PMF has a local minimum at *ξ*_0_, and a maximum at *ξ*_0_′ which is greater than *ξ*_0_.

Molecular structures were illustrated by using Visual Molecular Dynamics (VMD)^54^. Error bars were calculated from standard error that indicate 95% confidence interval.

## Results and Discussion

### Subcomplex species and disassembly pathway obtained from the hcbMC/MD simulations

We performed five independent hcbMC/MD simulations for the SAP system. A hcbMC/MD cycle is repeated 500 times, so that total simulation time length in the MD phase is 100 ns. The number of trial configuration M (*see* Materials and Methods for the definition) was set to 10 as an attempt; choice of M could be further optimized to enhance sampling. Nonetheless, the sampling efficiency of our method is similar to that of TRS method^20^ and seems sufficient to promote disassembly processes.

A ring shape SAP pentamer is converted into a set of subcomplexes and subunits by the 500^th^ cycle of hcbMC/MD simulations (Figure 2). We find a subunit pair with non-specific contacts, which were formed irrespective of inter-subunit interaction surfaces of SAP (*see* Figure 1B for comparison). Such non-specific inter-subunit reassociation results from a periodic simulation box whose size is insufficient for apparent spatial separation among five SAP subunit. Then we identified subcomplexes by excluding such a reassociated subunit pair that solely makes non-specific inter-subunit contacts.

**Figure 2.**
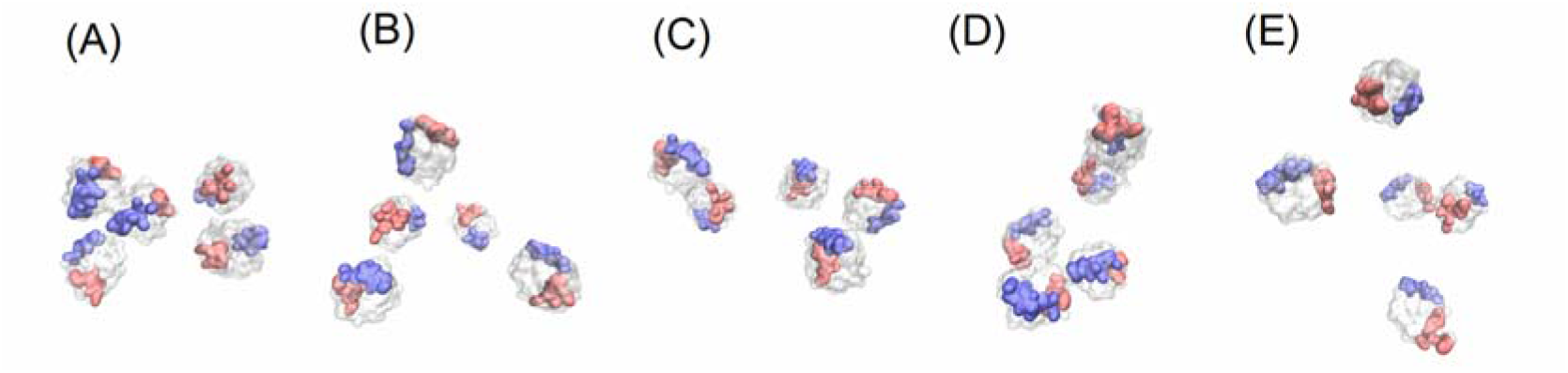
Configurations of SAP subunits at the last (500^th^) hcbMC/MD cycle, where each subunit pair has no native contacts, thus being supposed to be dissociated. Two individual inter-subunit interaction surfaces of each subunit are highlighted by blue and red colors as in the case of Figure 1. Alphabetic annotations for panels, (A)-(E), denote those for individual simulations hereafter.

Figure 3 shows subcomplex species at each hcbMC/MD cycle for the homo-pentameric SAP system. In our hcbMC/MD simulations, any of subcomplex species (tetramer, trimer and dimer) observed in the MS experiments^13^ appears. MS experiments induce inter-subunit dissociation reactions using atomic collision with gas molecules, while hcbMC/MD simulations accelerate these reactions with employing repulsive forces acting between a subunit pair. Although our hcbMC/MD scheme employs dissociation mechanisms differing from those in the MS experimental method^13^, our simulations might finely evaluate relative SAP-subunit interface strength similarly to the MS method.

**Figure 3.**
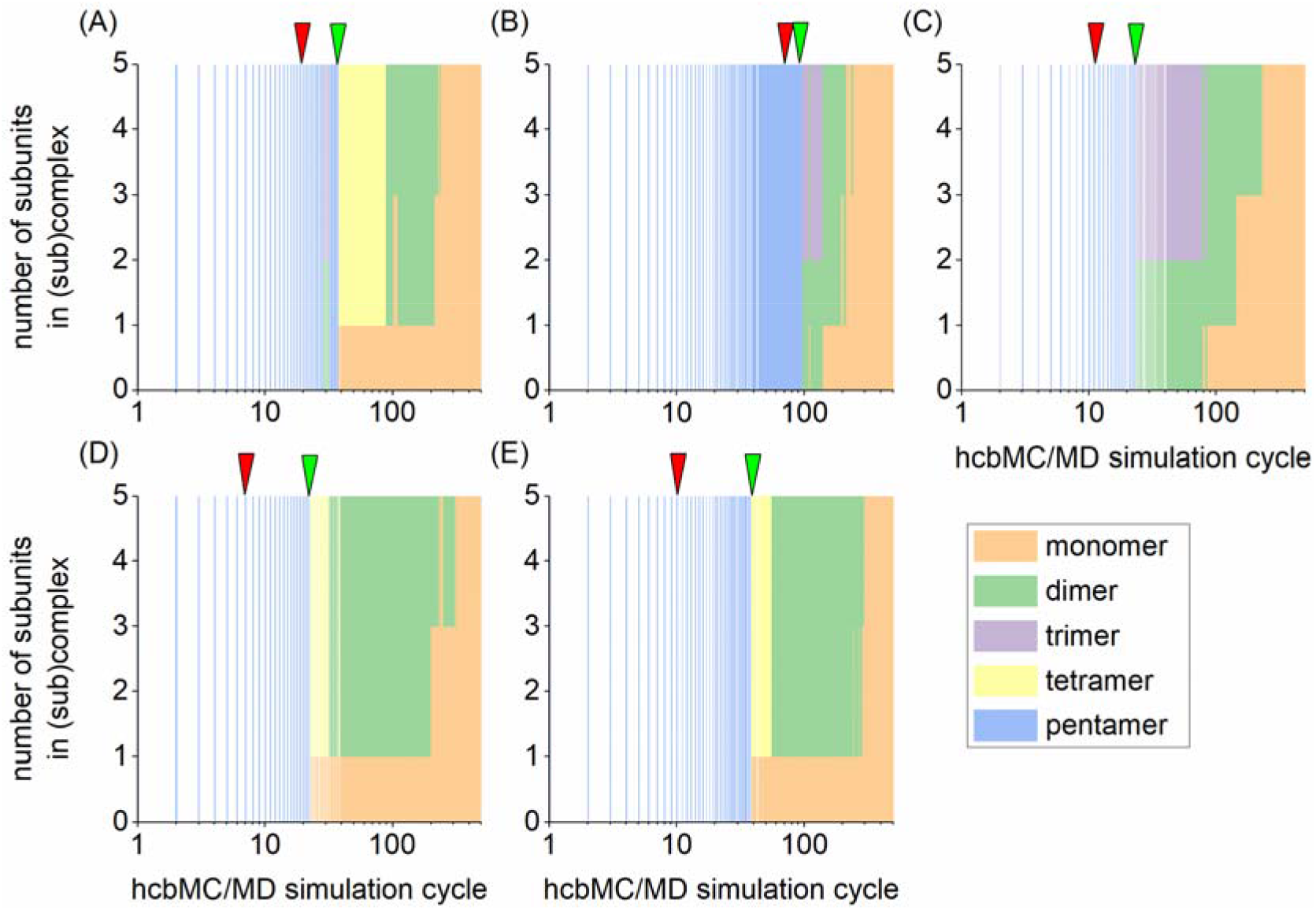
Disassembly of serum amyloid P component (SAP) homo-pentamer through hcbMC/MD cycles. Each panel is for an individual simulation. A distribution of SAP (sub)complex species is illustrated by color. Green and red arrows on panel indicate the initial stage of SAP disassembly (generation of tetramer or trimer) and the first cycle where SAP pentamer undergoes a ring-opening event.

Besides, each of the five simulations results in decomposition into the five isolated SAP subunits. Meanwhile the initial stage of SAP disassembly is classified into two distinct events: trimer-dimer formation and tetramer-monomer formation from the pentamer (Figure 3A, D and E, and Figure 3B and C, respectively; *see* Figure 1C). In Figure 3A, the trimer-dimer pair found at 29^th^ cycle returns to pentamer form, so that we suppose tetramer-monomer formation as the initial subcomplex disassembly. These two different disassembly events were similarly predicted by the structural bioinformatics approach in the earlier MS study.^13^

It is thus possible to say that the simulation results are consistent with those of the earlier study^13^ with respect to SAP subcomplex species and the initial stage of SAP disassembly processes. Our reaction pathway sampling method could play a role complementary to MS and SB approaches in arrangement of complete sets of atomic coordinates of all subunits involved in a disassembly process.

### Computation performance of hybrid configuration bias MC/MD simulations

The disassembly of pentameric form of SAP can be efficiently simulated due to methodological advantage of our hcbMC/MD approach. We can show this point by using independent five unbiased NPT MD simulations (300 K, 1 bar) with simulation length of 200 ns in total, twice longer than those of hcbMC/MD simulations. As shown in Figure 4, the five SAP molecules retain each inter-subunit contacts during the 200-ns simulations, thus keeping to take a pentameric form in each of the unbiased MD simulations.

**Figure 4.**
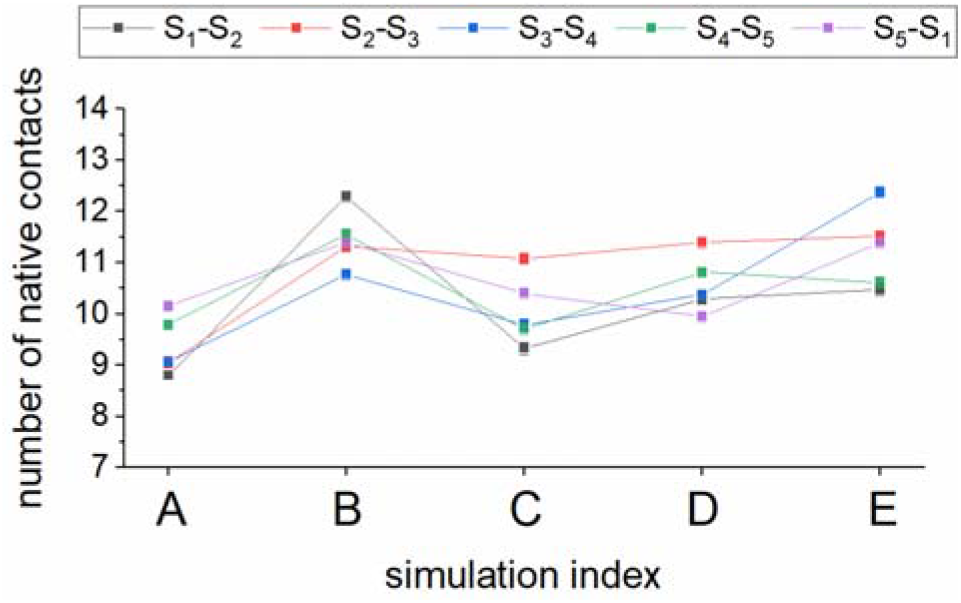
Averaged number of native contacts between SAP subunit pair. Average and statistical error values are calculated using 2000 snapshot structures obtained from each of the five unbiased 200-ns MD simulations (an error bar is buried in the square, then not being apparent in this panel). Each inter-subunit native contacts are distinguished by different color. Annotation of subunit pair is given as in the case of Figure 1. Statistical errors were estimated from standard error by considering 95 % confidence interval.

Furthermore, each of the subunits retains folded configurations though hcbMC/MD cycles. Although we accelerated dissociation reactions using external forces, the values of RMSd for SAP subunits are similar between hcbMC/MD simulations and unbiased 200-ns NPT MD simulations (**Table 1**). This observation clearly shows that external forces applied to subunits in SMD simulations hardly affect their structures, thus additionally supporting applicability of our hcbMC/MD method to analyze physicochemical properties of proteins under physiological conditions.

**Table 1.** Root mean square deviation (RMSd)^a^ with error value^b^ for SAP subunit^c^.

|  | S<sub>1</sub> | S<sub>2</sub> | S<sub>3</sub> | S<sub>4</sub> | S<sub>5</sub> |
| --- | --- | --- | --- | --- | --- |
| hcbMC/MD | 0.9 ± 0.2 | 0.9 ± 0.1 | 0.8 ± 0.1 | 0.8 ± 0.1 | 0.8 ± 0.1 |
| unbiased MD | 0.9 ± 0.2 | 0.8 ± 0.2 | 0.8 ± 0.2 | 0.8 ± 0.2 | 0.8 ± 0.3 |

<sup>a</sup>RMSd for each SAP subunit was averaged over each 500 hcbMC/MD cycles (100 ns in total) or 2000 unbiased MD cycle (200 ns in total), and averaged over the 5 simulations.
<sup>b</sup>Error values denote CI 95, derived from standard deviation over the 5 simulations.
<sup>c</sup>SAP subunit is distinguished by combination of alphabetic letter and digit as in the case of Figure 1.

### Free energy profile of ring-opening reaction of SAP pentamer

Observing the hcbMC/MD trajectories, we can find a preliminary dissociation event in each of the 5 simulations. One of the five subunit interaction interfaces is broken, then resulting in ring-opened pentameric form of SAP (Figure 5). A ring opening of the SAP pentamer occurs in a relatively early stage of hcbMC/MD cycles (Table 2) and is followd by the initial steps of SAP pentamer disassembly process, that is, tetramer-monomer and trimer-dimer dissociation reactions (*see* Figure 3).

**Figure 5.**
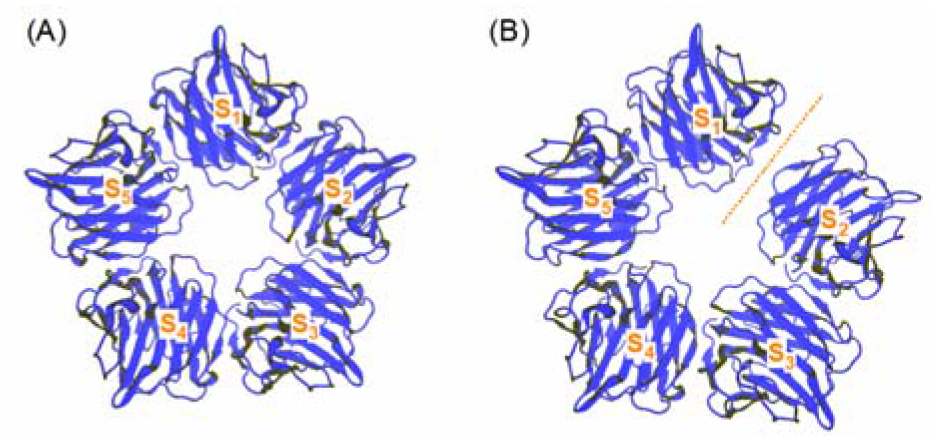
Ring-opened SAP pentamer configuration. (A) X-ray crystallographic structure (PDB entry: 4AVS). (B) Snapshot structure obtained from hcbMC/MD simulation, whose simulation index is A in Table 2. Orange dotted line is depicted in the vicinity of broken subunit interaction interface. SAP subunit is distinguished by combination of alphabetic letter and digit as in the case of Figure 1.

**Table 2.** Geometrical characterization of SAP pentamer at the cycles where the initial ring-opening reaction occurs.

| simulation index | pair of SAP subunit<sup>a</sup> | cycle | CCS<sup>b</sup> [nm<sup>2</sup>] |
| --- | --- | --- | --- |
| A | S<sub>3</sub>-S<sub>4</sub> | 20<sup>th</sup> | 59.9 |
| B | S<sub>1</sub>-S<sub>2</sub> | 61<sup>st</sup> | 59.8 |
| C | S<sub>3</sub>-S<sub>4</sub> | 11<sup>th</sup> | 59.1 |
| D | S<sub>2</sub>-S<sub>3</sub> | 7<sup>th</sup> | 59.3 |
| E | S<sub>3</sub>-S<sub>4</sub> | 10<sup>th</sup> | 59.8 |

<sup>a</sup>SAP subunit is distinguished by combination of alphabetic letter and digit as in the case of Figure 1.
<sup>b</sup>CCS denotes collision cross section.

This ring-opening reaction has not been reported in the earlier mass spectroscopy studies^13, 30, 31^, although there are rooms to discuss occurrence of the intermediate states under realistic experimental conditions. It is possible that MS observations have not detected such intermediate states due to their limitation for spatial resolution. Values of collision cross section (CCS) for these ring-opened SAP pentamer structures are around 59 nm^2^, being larger by c.a. 3 nm^2^ than that calculated from the X-ray crystallographically resolved ring-forming pentamer^32^. A ring-opened SAP pentamer appearing in experimental conditions might be assigned into partially unfolded SAP pentamer in the context of mass spectroscopy.^30^

We then consider possibility of experimental observation of this ring-opening reaction. High-speed timelapse atomic force microscopy (AFM) is now available to observe molecular dynamics of multimeric protein complexes with nanometer spatial resolution and hundreds of milliseconds temporal resolution.^55^ This method could be employed to validate the presence of this reaction, if the reaction time scale is found within observation period of AFM. Aiming to estimate the reaction time, we carried out free energy profile calculations with a conventional protocol, umbrella sampling molecular dynamics (USMD) simulations combined with Weighted Histogram Analyses Method (WHAM)^50, 51^.

We here examined the initial ring-opening events appearing in the hcbMC/MD simulations due to the following observation for the trajectories we obtained. Ring-opened SAP pentamer does not necessarily lead to immediate progress to the initial dissociation steps but may transiently come back to ring-forming intact SAP pentamer (*see* Figure 3 A and Figure S1 in Supporting Information). A SAP structure fluctuating between the two pentameric forms might gradually lose inter-SAP subunit contacts; such partially broken contacts would give lower activation barrier of the ring-opening reaction so that additional free energy calculations are probably needed to estimate a precedent free energy loss.

We then arranged a set of initial atomic coordinates for USMD simulations by using snapshot structures appearing at the cycle of initial ring-opening reaction and also those around the neighboring ones if needed. Each of these snapshot structures takes either ring-formed pentamer or ring-opened pentamer. A reaction coordinate is set to the distance between centers of gravity of subunits undergoing breakage of inter-subunit contacts (*see* Table 2 for subunit pairs examined for PMF calculations). We also calculated PMFs by using the initial structure for the hcbMC/MD simulations as references for stable SAP pentamer configuration, which are shown by grey lines in Figure 6A, 6B and 6C.

**Figure 6.**
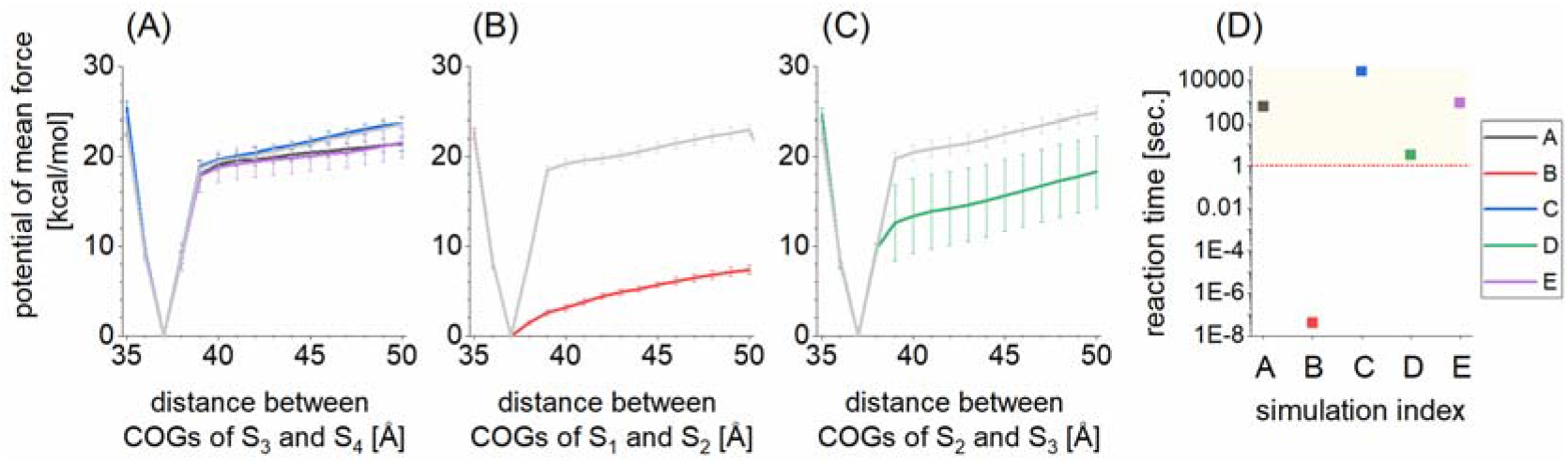
Potential of mean force (PMF) of SAP pentamer ring-opening reaction (A), (B) and (C), and reaction time evaluated with Eyring’s transition state theory (D). Each hcbMC/MD simulation is indexed by alphabetic character, A-E and distinguished from the remainings on panel by different color. An error bar was calculated from standard deviation and denoted by CI95. In panels (A), (B) and (C), PMFs calculated from the initial structure for the hcbMC/MD simulations are shown by grey lines. SAP subunit pair is distinguished by combination of alphabetic letter and digit as in the case of Figure 1. In panel (D), points above the red dotted borderline, netted with the orange square area, have the estimated reaction time falling within observation period with AFM experiments.

Each of PMFs has one activation barrier and appears to be up-hill (Figure 6A, 6B and 6C). A steep change of the slope of PMF is found between 37 and 40 Å on the reaction coordinates. Among the five hcbMC/MD derived-PMFs, the simulation indexed by C shows the highest activation barrier of 23.7 kcal/mol (*see* blue line in Figure 6A) and the corresponding reaction time is 8.2 hour according to eq. (2). This estimated reaction time may fall within observation periods of timelapse AFM, hundreds of milliseconds or longer, so that occurrence of this ring-opening reaction could be validated with employing AFM experiments. The above discussion leads to such a speculation that a part of SAP pentamer might have ring-opening forms during AMF observation period.

It is noted that the above speculation is similarly obtained even if we use Kramers’ transition state theory. Kramers’ theory possibly give longer reaction timescale than Eyring’s one^53^, although this does not essentially change the speculation; ring-opening reactions are still supposed to occur within AFM observation time scale.

Meanwhile, large deviations among the values of activation barrier might denote variations of ring-opening reaction pathways and also following disassembly processes perhaps. Recalling observation period of timelapse AFM experiments, we then classify the five ring-opening events into two groups; that for the simulation annotated by B (sim-B) and those for the other four. The reaction time obtained from sim-B is 39 ns, significantly smaller than those obtained from the other four simulations (Figure 6B). The reaction time of 39 ns means that a ring-opening reaction can spontaneously proceed under thermal fluctuation along this reaction pathway.

The significant decrease of activation barrier could be explained by difference in the number of atomic contacts between a subunit pair which retains contacts of inter-subunit interaction surfaces. Figure 7 shows the number of atomic contacts between each subunit pair for the five hcbMC/MD simulations, where the snapshot structures are obtained from the simulation cycle at which we observed the initial ring-opening reaction. With regard to inter-subunit atomic contacts retained in ring-opened SAP pentamer, we can find smaller number of atomic contacts in S_5_-S_1_ interface in sim-B, that is, 8. Meanwhile the remaining three are 15 or larger. It is of note that S_5_ is neighboring to S_1_, which undergoes ring-opening reaction in sim-B (Table 2). According to such a preliminary deformation at the vicinal interface, the reaction pathway obtained from the sim-B and those obtained from the other four might be separately located in the conformational space of this SAP system.

**Figure 7.**
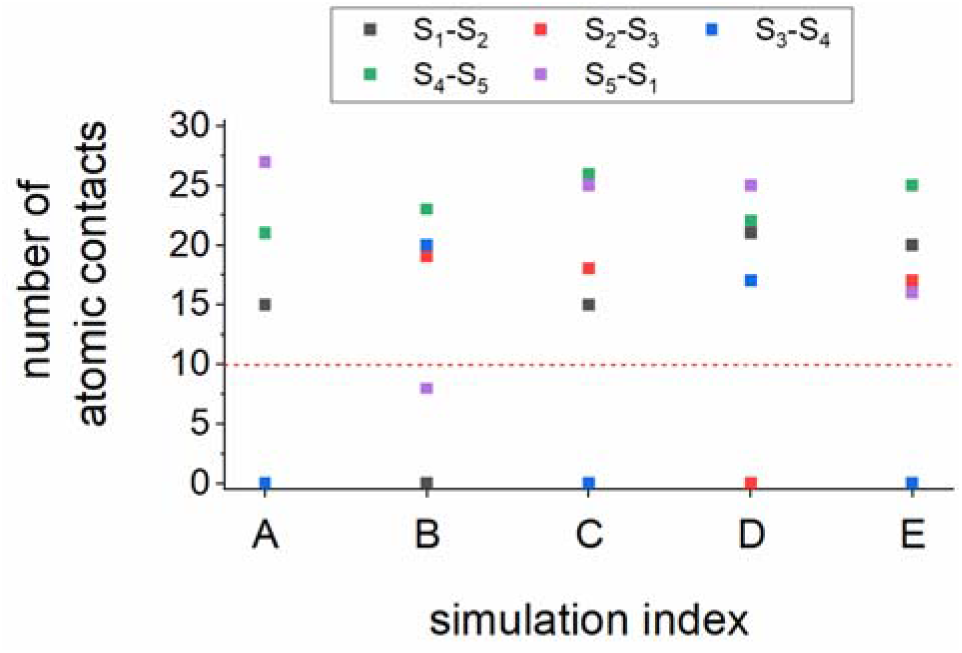
Atomic contacts between subunit pair for snapshot structure at hcbMC/MD cycle for the initial ring-opening reaction. SAP subunit is distinguished by combination of alphabetic letter and digit as in the case of Figure 1.

From the viewpoint of reaction energetics, stepwise breakage of subunit interface seems to more easily occur rather than simultaneous break of multiple subunit interfaces. Recalling the earlier simulation study on stepwise dissociation reaction of protein dimer^56^, we can assume that this ring-opening reaction proceeds in a similar manner; allosteric losing of inter-subunit contacts is followed by dissociation between a subunit pair.

We suppose that breakage of salt bridge formed between Lys117 and Asp42 in S_5_-S_1_ interface (Figure 8A, B and C) is particularly important to weaken configurational restraints between S_1_ and S_5_ in the ring-opened SAP pentamer in sim-B. This salt bridge breakage seems to allow libration motion of S_1_ on S_5_ and enhances dissociation of S_1_ from S_2_. The effects of Lys117 and Asp42 on SAP pentamer formation have not been reported to our best knowledge. While considering significant decrease of the activation barrier for sim-B, mutation on either or both of these two charged residues, or strongly acidic and basic conditions, might repress SAP oligomerization. SAP was identified as pathological deposite in 1967 and has been widely studied in the field of amyloid- and immune-related diseases.^57, 58^ These two residues might make a theraphetic target in aim of regulation of SAP pemtamer formation.

**Figure 8.**
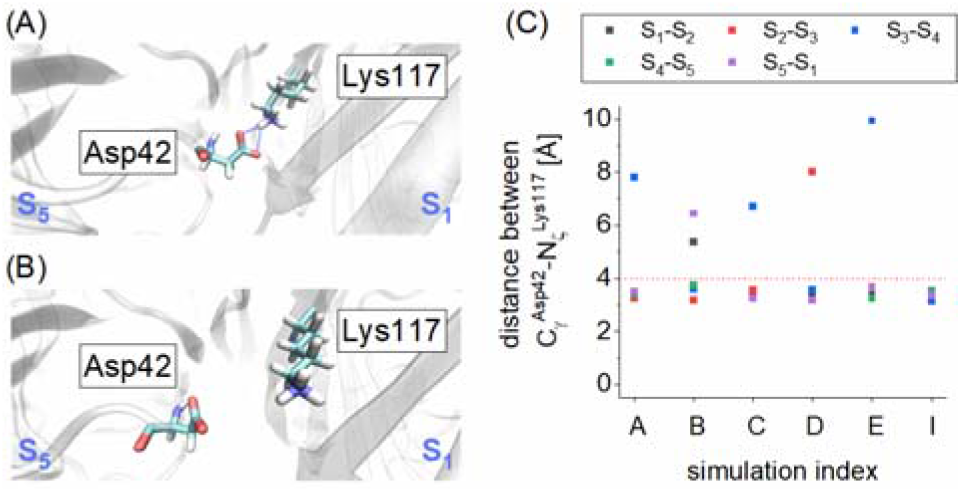
Positional relation between Asp42 and Lys117 at subunit interface. (A) The initial structure of hcbMC/MD simulations. (B) Structure at the timing of initial ring-opening in the simulation indexed with B. (C) Distances between Cγ in Asp42 and Nζ in Lys117 for each interface, where simulation index I denotes the initial structure of hcbMC/MD simulations. SAP subunit is distinguished by combination of alphabetic letter and digit as in the case of Figure 1. In Panel A, blue dotted lines denote hydrogen bonding between Lys117 and Asp42. In Panel C, red dotted line shows level of 4 Å.

According to the above discussion, we can speculate that atomistic events occurring in the hcbMC/MD simulation, such as preliminary deformation at vicinal SAP subunit interface, has influence on activation barrier of the ring-opening reaction. This speculation is supported from comparison between the two free energy profiles; one for ring-opening simulated from an hcbMC/MD trajectory, other for ring-opening simulated with the initial structure of hcbMC/MD, that is, reference to stable SAP pentamer configuration. We can find substantial free energy decrease of 11 kcal/mol in Sim-B and that of 6 kcal/mol in Sim-D at 50 Å on each reaction coordinate (*see* Figure 6B and C, respectively).

In particular, compared with the other four simulations, the change for Sim-B is so large as to make the timescale of the ring-opening reaction 10^7^-fold smaller than time resolution of AFM observations (*see* Figure 6D). The above observation indicates physicochemical importance of the preliminary deformation. The ring-opening event observed in Sim-B might be essentially a two-step reaction accompanying the preliminary deformation. Besides, the preliminary deformation itself is possibly a time-consuming process, which is characterized by free energy profile with barrier of a substantial height, thus being the rate-determining step of ring-opening reaction pathway observed in Sim-B.

We here have discussed the reaction mechanism in terms of one reaction coordinate, distance between centers of gravity of a subunit pair involved in interface breakage. We could refine physicochemical characterization of the reaction mechanism by employning an additional reaction coordinate, *e.g*., the number of allositeric inter-subunit contacts at the vicinal interface. Similarly, time scale of preliminary deformation could also be elucidated quantitatively and the actual reaction time of ring opening pathway in Sim-B could be obtained. Nontheless, further efforts to address these problems are beyond research scope of this study, thus being left for future studies.

## Concluding Remarks

We have here proposed a new reaction pathway sampling method, hcbMC/MD, aiming to simulate complicated assembly and disassembly processes involving multimeric biomacromolecules within realistic computation time.

Our method was applied to disassembly simulations of Serum Amyloid P component (SAP) homo pentamer. Subcomplex species and disassembly pathway we obtained from hcbMC/MD simulations are consistent with the earlier observations. Our simulation approach could play a role complementary to mass spectrometry and structural bioinformatics approaches with regard to arrangement of complete sets of atomic coordinates of a molecular system undergoing disassembly processes.

Regarding observations obtained from mass spectroscopy and structural bioinformatics approaches as molecular and structural landmarks, our atomistic simulation approaches could provide deeper insights into physicochemical mechanism for biomacromolecular assembly and disassembly processes. Actually we observed a novel dissociation event, ring-opening reaction of SAP pentamer. Employing free energy calculation combined with our hcbMC/MD trajectories, we moreover obtained experimentally testable observations on (1) reaction time of the ring-opening reaction and (2) importance of Asp42 and Lys117 for stable formation of SAP oligomer.

Considering physicochemical complexity of atomistic multicomponent systems and practical restriction of computational resources, a configuration ensemble generated by any enhanced sampling method is probably a subset of an exact configurational space. Although this viewpoint often raises such a concern that enhanced sampling simulations for biomacromolecules pick up biologically insignificant configurations, we suppose that our hcbMC/MD method is practically well-designed to extensively sample multimeric biomacromolecule disassembly pathway and examine the mechanisms at atomic level (an additional remark on our hcbMC/MD derived configuration ensemble is given in Supporting Information).

As shown above, our pathway sampling simulations succeeded to convert SAP pentamer into the five SAP subunits within realistic computational time. SAP subcomplex species appearing through the disassembly processes were consistent with those deduced from the MS experiments. Furthermore, we obtained experimentally testable predictions for the disassembly processes and mechanisms via free energy calculations. We then suppose that our hcbMC/MD approach opens a new avenue to study physicochemical mechanisms of complicated biological processes associated with multimeric biomacromolecules in the cell.

## Supporting Information

See supporting information for detailed procedures for unbiased MD, SMD and USMD simulations.

## Acknowledgements

This work was supported by a Grant-Aid for Scientific Research on Innovative Areas “Chemistry for Multimolecular Crowding Biosystems” (JSPS KAKENHI Grand No. JP17H06351).

## TOC graphics

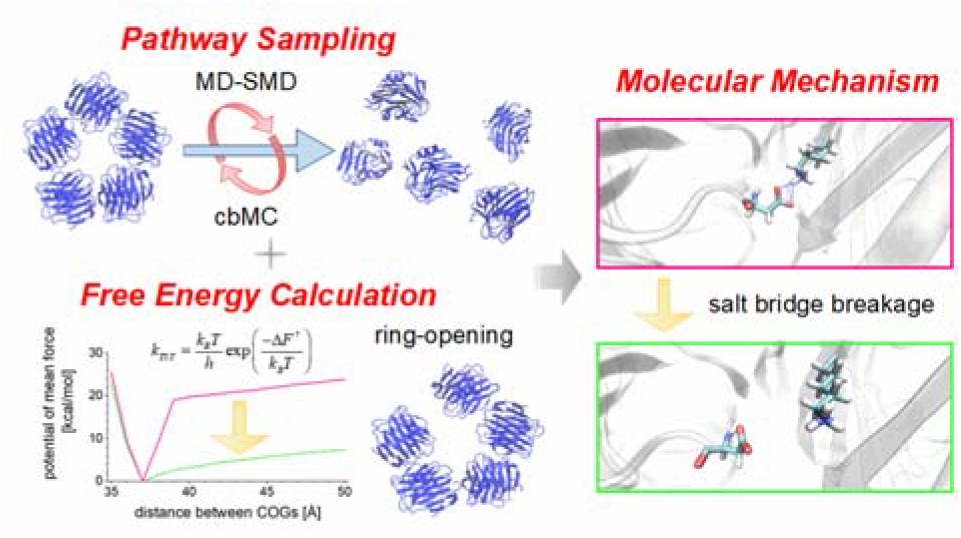

