## Supporting Information for "Reaction Pathway Sampling and Free Energy Analyses for Multimeric Protein Complex Disassembly with Employing Hybrid Configuration Bias Monte Carlo/Molecular Dynamics Simulation"

### S-1. Setup of molecular dynamics simulations

In each simulation, electrostatic interaction was treated by the Particle Mesh Ewald method, where the real space cutoff was set to 9 Å. The vibrational motions associated with hydrogen atoms were frozen by SHAKE algorithm through MD simulations. The translational center-of-mass motion of the whole system was removed by every 500 steps to keep the whole system around the origin, avoiding an overflow of coordinate information from the MD trajectory format. Snapshot structures were recorded by every 10 ps. These simulation conditions referred above were common in all of the simulations discussed in this manuscript.

### S-2. Preparation of initial atomic coordinates for hcbMC/MD simulations

Each atomic coordinate of water and $Cl^-$ molecules in the SAP system was energetically relaxed by the following molecular mechanics (MM) and molecular dynamics (MD) simulations. First, steric clashes in the system were removed by MM simulation, which consists of 1000 steps of the steepest descent method followed by 49000 steps of the conjugate gradient method.

Then the system temperature and density were relaxed through the following five MD simulations: NVT (0.001 to 1 K, 0.1 ps) → NVT (1 K, 0.1 ps) → NVT (1 to 300 K, 20 ps) → NVT (300 K, 20 ps) → NPT (300 K, 300 ps, 1 bar). In each of MM and MD simulations, the atomic coordinates of non-hydrogen atoms in SAP were restrained by the harmonic potential with force constant of 10 kcal/mol/$Å^2$ around the initial atomic coordinates. The first two NVT MD simulations and the other MD simulations were performed using 0.01 fs and 2 fs for the time step of integration, respectively. The first and second NVT MD simulations were performed using Berendsen thermostat[1] with a 0.001 ps of coupling constant. Meanwhile the following three simulations were performed using Langevin thermostat with 1-$ps^{-1}$ of collision coefficient. In the first NVT MD simulation, the reference temperature was linearly increased along the time-course.

In the NPT MD simulation, the system pressure was regulated with Monte Carlo barostat, where the system volume change was attempted by every 100 steps. Each set of initial atomic velocities was randomly assigned from the Maxwellian distribution at 0.001 K.

For each atomic coordinates obtained above, a SAP conformation also was structurally relaxed in aqueous solution through the following 7-step MD simulations: NVT (0.001 to 1 K, 0.1 ps, 10 kcal/mol/Å) → NVT (1 to 300 K, 0.1 ps, 10 kcal/mol/Å) → NVT (300 K, 10 ps, 10 kcal/mol/Å) → NVT (300 K, 40 ps, 5 kcal/mol/Å) → NVT (300 K, 40 ps, 1 kcal/mol/Å) → NVT (300 K, 40 ps) → NPT (300 K, 1 bar, 10 ns). The first two NVT MD simulations and the other MD simulations were performed using 0.01 fs and 2 fs for the time step of integration, respectively. In the first two NVT simulations, the reference temperature was linearly increased along the time-course. In the first 5 steps, non-hydrogen atoms in SAP were positionally restrained by the harmonic potential around the initial atomic coordinates. In each NVT simulation, temperature was regulated using Langevin thermostat with 1-ps$^{-1}$ collision coefficient. In the last 10-ns NPT simulation, temperature and pressure were regulated by Berendsen thermostat[1] with a 5-ps coupling constant, and Monte Carlo barostat, where system volume change was attempted by every 100 steps, respectively. The initial atomic velocities were randomly assigned from the Maxwellian distribution at 0.001 K. The snapshot structure obtained from the 10-ns NPT

MD simulation procedure was employed for the following hcbMC/MD simulations.

### S-3. Configuration generation by using unbiased MD and steered MD simulations

The system temperature and pressure were regulated by Langevin thermostat with a 1-ps collision coefficient, and Monte Carlo barostat with attempt of system volume change by every 100 steps, respectively. The trajectory was recorded every 10-ps interval. In each of MD and SMD simulations, a set of initial atomic velocities was calculated from products of integration time step and atomic forces acting on the atoms.

A corresponding unbiased NPT-MD simulation consists of 2000 cycle of 100-ps unbiased MD simulation. A set of atomic velocities was reset in each cycle similarly to the above hcbMC/MD simulations. Remaining simulation conditions are similar to those for the hcbMC/MD simulations.

### S-4. Umbrella sampling molecular dynamics simulations

Using each initial atomic coordinates derived from the hcbMC/MD simulations, we performed the relaxation simulation and the following USMD simulation. The relaxation simulation consists of 5 steps: NVT (0.001 to 1.0 K, 0.1 ps, 100 kcal/mol/Å$^2$) → NVT (1.0 to 300 K, 0.1 ps, 100 kcal/mol/Å$^2$) → NVT (300 K, 40 ps, 10 kcal/mol/Å$^2$) → NVT (300 K, 40 ps, 5 kcal/mol/Å$^2$) → NVT (300 K, 40 ps, 1 kcal/mol/Å$^2$). The initial atomic velocities were randomly assigned from the Maxwellian distribution at 0.001 K in the first step. Backbone heavy atoms (C$\alpha$, C, N, O) of SAP pentamer were restrained by the harmonic potential around the initial atomic coordinates. In each of the first two NVT-MD simulations, the reference temperature was linearly increased along the time-course. Then, several nanosecond USMD simulation was executed under NVT condition (300 K). For each system, the USMD simulation time length is determined by evaluating the convergence of PMF (*see* Figure S2 and Table S1). The reaction coordinate of these USMD simulations is the distance between centers of gravity of subunits undergoing full dissipation of atomic contacts. Minimum of the harmonic potential and the corresponding force constant for window of each USMD simulation are summarized in Table S1. In an USMD simulation, temperature was regulated using Langevin thermostat with 1-ps$^{-1}$ collision coefficient, and the interatomic distance for reaction coordinate was recorded every 1-ps interval.

Using 8 sets of USMD simulations, we made a complete histogram spanning the reaction coordinate from 33 to 53. We confirmed 3.5% or greater overlap between sampling of neighboring USMD windows, indicating accurate construction of potential of mean force by WHAM[2, 3]. Then the complete histogram was employed to compute a potential of mean force, where width of bin was set to 1 Å.

### S-5. Steered molecular dynamics-combined umbrella sampling molecular dynamics simulations

Steered molecular dynamics (SMD) simulations, starting from the initial atomic coordinates for the hcbMC/MD simulations, were performed to prepare the initial atomic coordinates for each window of umbrella sampling (US) MD simulations.

As the reaction coordinate for SAP ring opening, we consider the distance between centers of gravity of C$\alpha$ atoms in each of SAP subunit pair. Here we employ '$S_1$ and $S_2$', '$S_2$ and $S_3$' and '$S_3$ and $S_4$' as a reaction pair, because the ring opening reactions associated with these three pairs are discussed in the main text (*see* Figure 6A-C).

The value of reaction coordinate was gradually changed through the SMD simulations by imposing the harmonic potential with force constant of 10 kcal/mol/Å$^2$. The target distance of SMD simulations was set to 65 Å. The SMD simulation was executed for 0.25 ns under NPT condition (300 K; 1 bar). The trajectory was recorded by every 5 ps. Temperature and pressure were regulated using Langevin thermostat with a 1-ps$^{-1}$ of collision coefficient and Berendsen barostat [1] with a 5-ps coupling constant, respectively. A time step of 2 fs was used to integrate Newton's equation of motion. Initial atomic velocities were taken over from the previous MD simulation. Using each SMD trajectory, we prepared 16 snapshot structures for 16 umbrella windows (Table S2), where a value of the reaction coordinate for each snapshot structure ranges from 35 Å to 50 Å with interval of 1 Å. For each of the three reaction coordinates (*see* Figure 6A-C), this procedure was repeated 8 times and, the derived snapshot structures were employed for the following USMD simulations.

Using each initial atomic coordinates derived from the SMD simulations, we performed the relaxation simulation and the following USMD simulation. The relaxation simulation consists of 5 steps: NVT (0.001 to 1.0 K, 0.1 ps, 100 kcal/mol/Å$^2$) → NVT (1.0 to 300 K, 0.1 ps, 100 kcal/mol/Å$^2$) → NVT (300 K, 40 ps, 10 kcal/mol/Å$^2$) → NVT (300 K, 40 ps, 5 kcal/mol/Å$^2$) → NVT (300 K, 40 ps, 1 kcal/mol/Å$^2$). The initial atomic velocities were randomly assigned from the Maxwellian distribution at 0.001 K in the first step. Backbone heavy atoms (Cα, C, N, O) of SAP pentamer were restrained by the harmonic potential around the initial atomic coordinates. In each of the first two NVT-MD simulations, the reference temperature was linearly increased along the time-course. Then, 5-ns USMD simulation was executed under NVT condition (300 K). For each system, the USMD simulation time length is determined by evaluating the convergence of PMF (*see* Panels A-C in Figures S3). The reaction coordinate of these USMD simulations is the same as that for the SMD simulations. Minimum of the harmonic potential and the corresponding force constant for window of each USMD simulation are summarized in Table S2. In an USMD simulation, temperature was regulated using Langevin thermostat with 1-ps$^{-1}$ collision coefficient, and the interatomic distance for reaction coordinate was recorded every 1-ps interval.

Using 8 sets of 16 USMD simulations, we made a complete histogram spanning the reaction coordinate from 35 to 50 Å. We confirmed 2.5% or greater overlap between samplings of neighboring USMD windows, indicating accurate construction of potential of mean force by WHAM[2, 3]. Then the complete histogram was employed to compute a potential of mean force, where width of bin was set to 1 Å.

### S-6. Remarks on multi-trajectory approaches as potentially applicable methods

Multi-trajectory approaches, *e.g.*, Weighted Ensemble (WE) method[4], could be potentially applicable to solve problems we are addressing in this study. Since the WE method has already been employed to study an elementary process of multimeric subunit assembly, protein-peptide dimer formation at atomic level[5], applications to multimeric subunit assembly and disassembly processes appear to be a simple extension of this method.

Nonetheless, it still seems to be challenging for atomistic WE simulations to examine assembly/disassembly processes of multimeric biomacromolecule complexes with available computational resources of today. Even if they work, it probably costs substantially huge amount of computation time to simulate a biomacromolecule system with multiple subunit interfaces.

This observation is obtained by recalling the technical design of multi-trajectory approaches, dependence of sampling performance on the replica number and time length of unbiased MD simulation run. As for the recent atomistic WE simulations, length of each replicated MD trajectory is < 100 ps[5]. Relatively short sampling time raises a concern for magnitude of configurational displacement in each replicated trajectory.

Estimating translational diffusion coefficients of SAP from a protein similar in size, 20~30 kDa[6, 7], the value is 100 μm$^2$/s and expected spatial displacement within 100 ps is 1 Å at most; actual displacement should be further suppressed by crowding effect under inter-subunit interaction. Actually, employing displacement of distance between centers of gravity of SAP subunits during 100 ps as a measure to configuration displacement of SAP pair, a set of the values, calculated from each of the five 200-ns unbiased MD simulations of the SAP pentamer system, shows the average and standard deviation around 0.0 and 0.45 Å, respectively (Figure S4A). Positive and negative displacements seem to occur similarly thus being randomly (Figure S4B-F).

Due to such relatively small magnitude and randomness of the displacement, gain/loss of atomic contacts in one subunit interface would be recovered while other subunit pair gains/loses their atomic contacts, then retarding progress of assembly/disassembly of multimeric complexes; it thus seems to be difficult to obtain and keep substantial changes at each interface simultaneously.

This observation could similarly hold to other multi-trajectory approaches employing unbiased MD simulations, meanwhile indicating practical advantage of our hcbMC/MD method.

### S-7. Remark on configurations obtained from hybrid MC/MD simulations

A configuration ensemble generated by any enhanced sampling method is probably a subset of an exact configurational space. This observation is illustrated in Figure S5. Recalling this illustration, we give a remark of configuration space we sampled in this study and future perspective to developing our hcbMC/MD method.

We employed the hcbMC/MD simulations to accelerate inter-subunit dissociation reactions, which is challenge to access with brute-force MD simulations (*see* orange crosses and red arrow in Figure S5). Besides, the total time length of hcbMC/MD simulation is 100 ns at maximum, which is longer by 1000 times at maximum than lifetimes of salt-bridge formation discussed in some earlier studies[8, 9]. It then seems physicochemically reasonable that we could observe preliminary deformation (black cross in Figure S5), that is, breaking of salt bridge between Asp42 and Lys117 through a hcbMC/MD simulation.

Meanwhile we did not explicitly accelerate other slower reactions, which are relatively fast compared with disassembly processes but cannot be approached by using brute-force MD, for example, partial unfolding of monomeric SAP subunit (purple cross in Figure S5). Thus such processes would not be necessarily sampled with employing the current setup of hcbMC/MD simulations. Subtle RMSd values and the deviation, shown in Table 1, indicate that each of SAP subunits retains a folded conformation through hcbMC/MD simulations.

Nonetheless this point is less important as for our study on the SAP pentamer system. Coupling between (un)binding and unfolding is often focused upon studies on intrinsic disorder proteins (IDP),[10-12] although we confirmed by using the latest neural network-based IDP prediction methods[13, 14] that a SAP subunit has no disorder region except for short segments at N- and C-terminals (Figure S6), thus being supposed to be the ordered proteins rather IDPs. Then we suppose that SAP configurations sampled by the hdcMC/MD simulations are practically sufficient as initial guess of disassembly pathway of SAP pentamer.

**Table S1.** Equilibrium positions of biased potentials and corresponding force constants for subunit pair-dissociation umbrella sampling molecular dynamics simulations with hcbMC/MD simulation-derived configurations.

| equilibrium position of biasing potential [Å] | force constant [kcal/mol/$Å^2$] | | |
|---|---|---|---|
| | 10 | 50 | 100 |
| 35 | ✓ | | |
| 36 | ✓ | | |
| 37 | ✓ | | |
| 38 | ✓ | ✓ | ✓ |
| 39 | ✓ | ✓ | ✓ |
| 40 | ✓ | ✓ | ✓ |
| 41 | ✓ | ✓ | |
| 42 | ✓ | ✓ | |
| 43 | ✓ | | |
| 44 | ✓ | | |
| 45 | ✓ | | |
| 46 | ✓ | | |
| 47 | ✓ | | |
| 48 | ✓ | | |
| 49 | ✓ | | |
| 50 | ✓ | | |

**Table S2.** Equilibrium positions of biased potentials and corresponding force constants for subunit pair-dissociation umbrella sampling molecular dynamics simulations with the initial configuration for hcbMC/MD simulations.

| equilibrium position of biasing potential [Å] | force constant [kcal/mol/Å$^2$] | | | |
|---|---|---|---|---|
| | 10 | 50 | 100 | 150 |
| 35 | ✓ | | | |
| 36 | ✓ | | | |
| 37 | ✓ | | | |
| 38 | ✓ | ✓ | ✓ | |
| 39 | ✓ | ✓ | ✓ | ✓ |
| 40 | ✓ | ✓ | ✓ | ✓ |
| 41 | ✓ | ✓ | | |
| 42 | ✓ | ✓ | | |
| 43 | ✓ | | | |
| 44 | ✓ | | | |
| 45 | ✓ | | | |
| 46 | ✓ | | | |
| 47 | ✓ | | | |
| 48 | ✓ | | | |
| 49 | ✓ | | | |
| 50 | ✓ | | | |

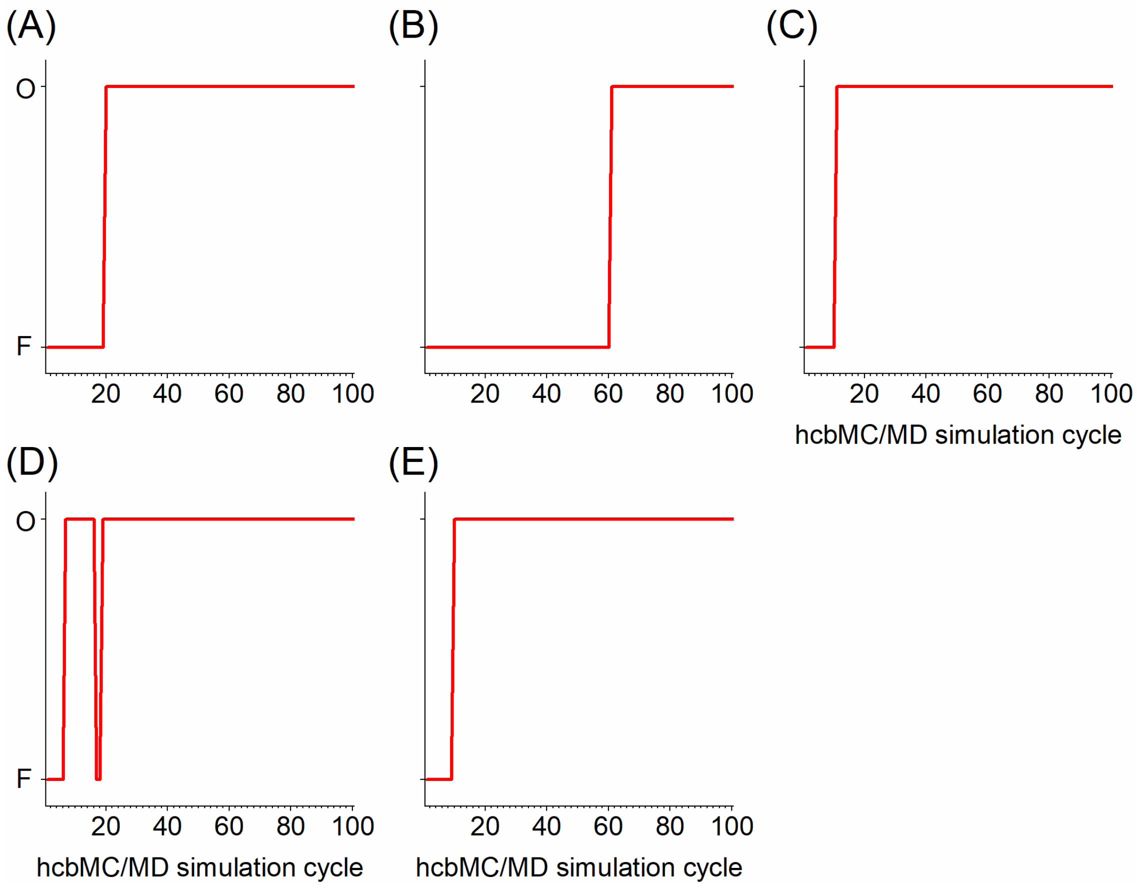


**Figure S1.** Monitoring pentameric form of SAP by the first 100 hcbMC/MD cycles. F and O beside longitudinal axes denote ring formation and ring opening conditions, respectively.

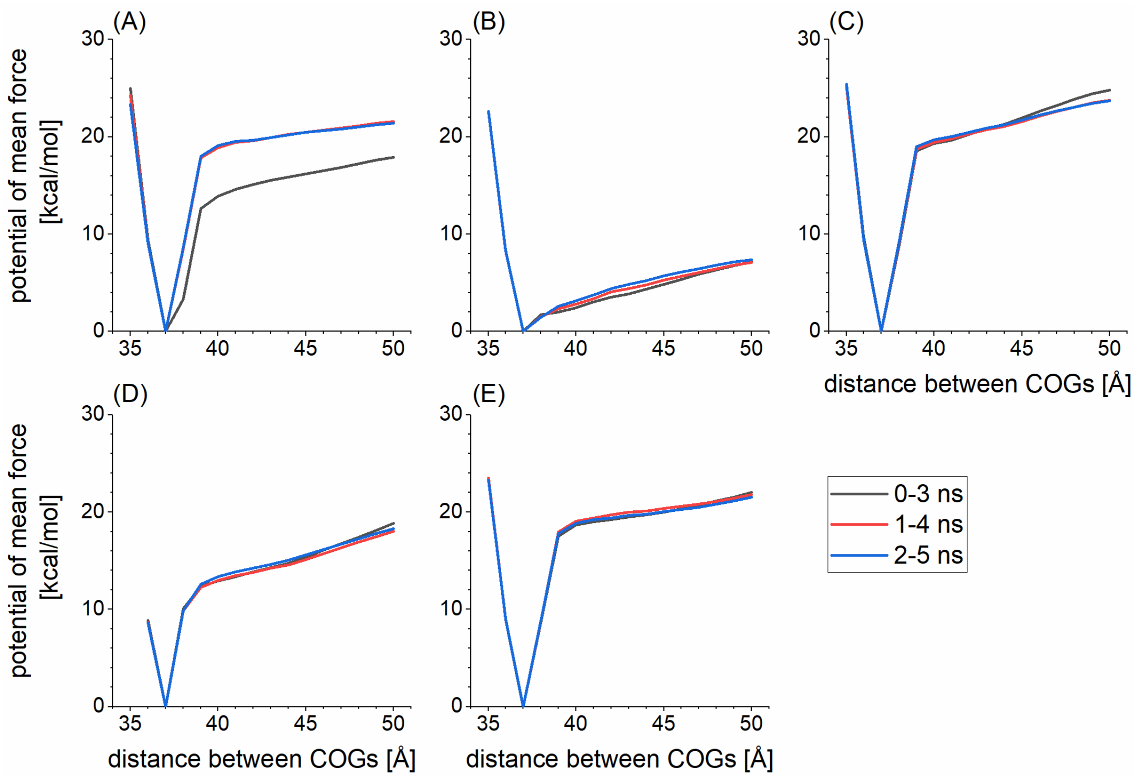


**Figure S2**. Convergence of potential of mean force of ring-opening reaction, calculated by using hcbMC/MD trajectories. Alphabetic annotations for panels corresponds to those for individual simulations discussed in the main text.

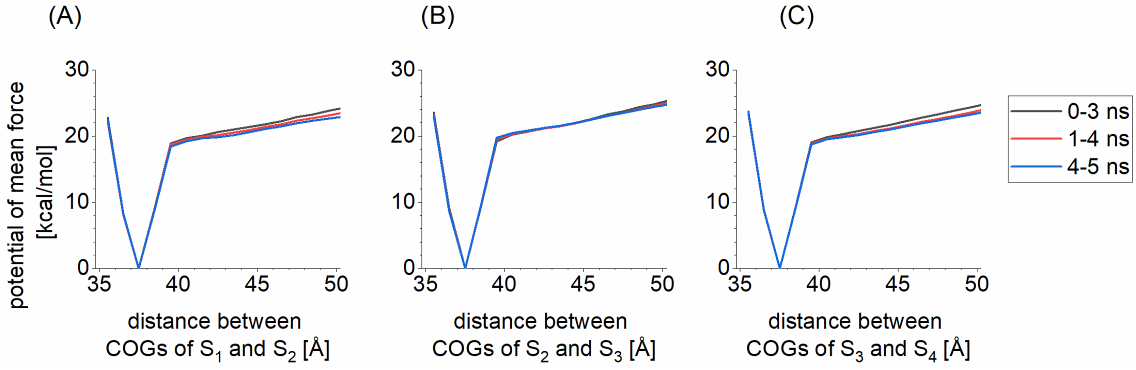


**Figure S3**. Convergence of potential of mean force of ring-opening reaction, calculated from the initial structure for the hcbMD/MD simulations. Alphabetic annotations for panels correspond to those for individual simulations discussed in the main text. SAP subunit pair is distinguished by combination of alphabetic letter and digit as in the case of Figure 1.

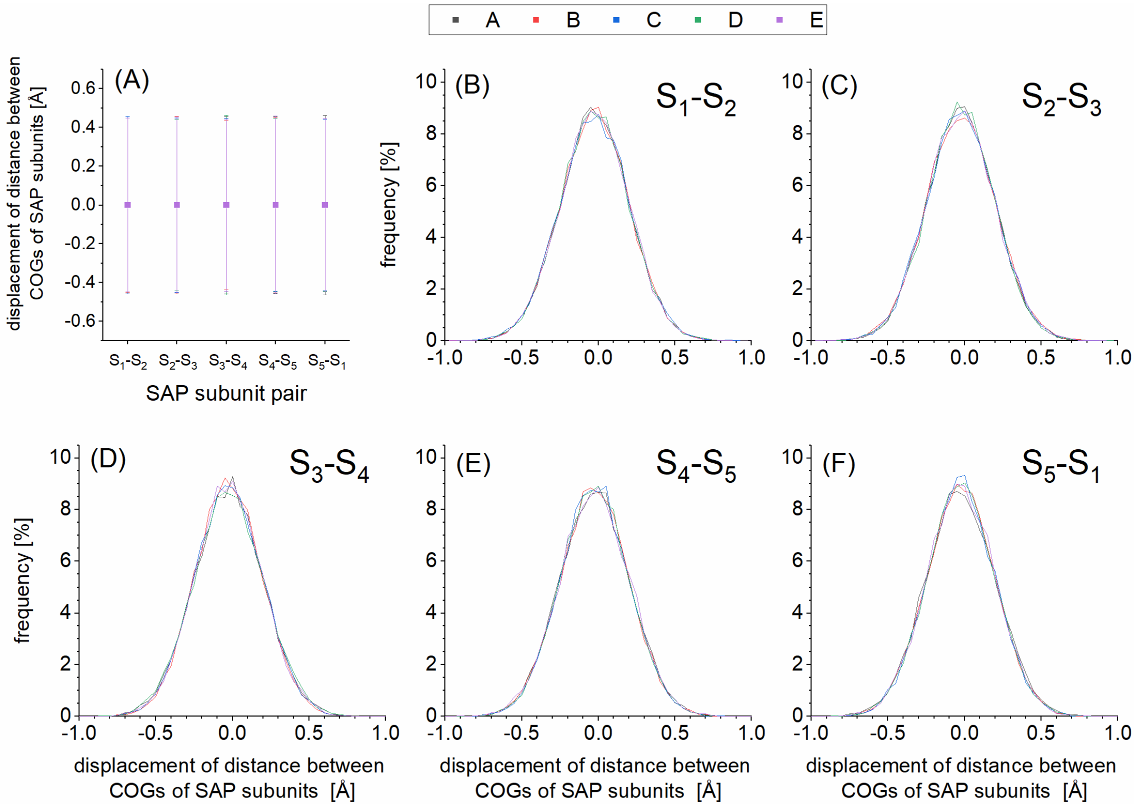


**Figure S4.** Displacement of distance between centers of gravity (COG) of SAP subunits during 100-ps period. (A) averaged value with error bar; (B)-(F) distribution of values for each subunit pair. Each hcbMD/MD simulation is indexed by alphabetic character, that is, A-E and distinguished from the remainings on panel by different color. SAP subunit is annotated by combination of alphabetic letter and digit as in the case of Figure 1. In panel A, error bar indicates 95% confidence interval, where an error is estimated with standard deviation. In panels B-F, a subunit pair is annotated at the upper right.

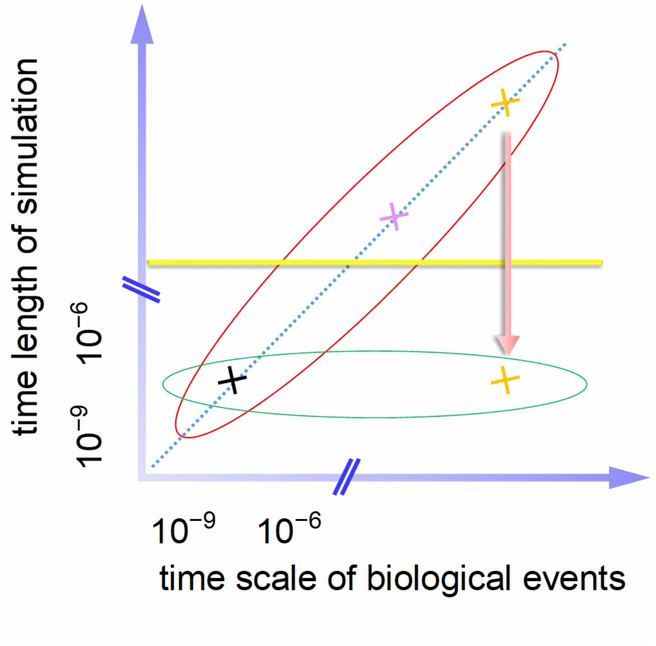


**Figure S5.** Conceptual illustration for time scale of biological events considered in configuration ensemble obtained from rare event sampling simulation. Red open and green open circles represent configuration ensembles by using brute-force simulations (BF) and rare-event sampling simulations (RS), respectively. Black cross denotes faster biological events (*e.g.*, reorientation of amino acid side chain) by both BF and RS. Orange and purple crosses denote slow biological events inaccessible by BF dynamics (e.g., folding of huge protein, disassembly of multimeric protein complex). Red arrow shows acceleration of the slowest dynamics here, represented by the orange cross.

```
                     •••••         •• •••        ••••••••••       ••••
              1 HTDLSGKVFV FPRESVTDHV NLITPLEKPL QNFTLCFRAY SDLSRAYSLF  50
SPOT            xxx
NetSurfP        xxx

                ••••       ••••••• •• •••••• ••• ••••        •••••••••
             51 SYNTQGRDNE LLVYKERVGE YSLYIGRHKV TSKVIEKFPA PVHICVSWES 100
SPOT
NetSurfP

                  ••••••••  ••••••••           • •••            •••••
            101 SSGIAEFWIN GTPLVKKGLR QGYFVEAQPK IVLGQEQDSY GGKFDRSQSF 150
SPOT
NetSurfP

                 •••••••••     •••••• ••••         •••    • •••   ••••
            151 VGEIGDLYMW DSVLPPENIL SAYQGTPLPA NILDWQALNY EIRGYVIIKP 200
SPOT
NetSurfP

                •
            201 LVWV 204
SPOT
NetSurfP          xx
```

**Figure S6.** Prediction of intrinsic disorder region of SAP subunit. 'SPOT' and 'NewSurfP' denotes SPOT-Disorder-Single [14] and NetSurfP-2.0 [13], respectively. Residues assigned to disorder are shown by cross. Closed circles over one-letter annotations of amino acid residue denote secondary structure; orange and green for α-helix and β-strand, respectively. Amino acid residues forming hydrogen bond and salt bridge between SAP subunits are highlighted by coloring in red.

6. Ferrage, F.; Zoonens, M.; Warschawski, D. E.; Popot, J. L.; Bodenhausen, G.,

Slow diffusion of macromolecular assemblies by a new pulsed field gradient NMR method. *J. Am. Chem. Soc.* **2003**, *125*, 2541-2545.

12. Arai, M.; Sugase, K.; Dyson, H. J.; Wright, P. E., Conformational propensities

of intrinsically disordered proteins influence the mechanism of binding and folding. *P. Natl. Acad. Sci. USA* **2015**, *112*, 9614-9619.

13. Klausen, M. S.; Jespersen, M. C.; Nielsen, H.; Jensen, K. K.; Jurtz, V. I.; Sonderby, C. K.; Sommer, M. O. A.; Winther, O.; Nielsen, M.; Petersen, B.; Marcatili, P., NetSurfP-2.0: Improved prediction of protein structural features by integrated deep learning. *Proteins* **2019**, *87*, 520-527.

14. Hanson, J.; Paliwal, K.; Zhou, Y., Accurate Single-Sequence Prediction of Protein Intrinsic Disorder by an Ensemble of Deep Recurrent and Convolutional Architectures. *J. Chem. Inf. Model.* **2018**, *58*, 2369-2376.
